# Endothelial Caveolin-1 and CXCL10 promote transcellular migration of autoreactive T cells across the blood-brain barrier

**DOI:** 10.1101/2022.11.15.516689

**Authors:** Troy N. Trevino, Ali A. Almousawi, Andrea Ochoa-Raya, Kait Zemanski, Suellen DS Oliveira, Felecia M. Marottoli, Leon M. Tai, Richard D. Minshall, Sarah E. Lutz

## Abstract

CXCL10 is an interferon-inducible chemokine that can recruit CXCR3^+^ leukocytes to the central nervous system, leading to neuroinflammation, demyelination, and neuronal losses. How CXCL10 promotes leukocyte extravasation and diapedesis across the blood-brain barrier – formed by brain endothelial cells – is poorly understood. Here, we report that CXCL10 mediates CD4+ T cell migration through the brain endothelial cell cytoplasm (transcellular), but not cell-cell junctions (paracellular), via the vesicular trafficking protein Caveolin-1. Caveolin-1 promotes CXCL10 aggregation into cytoplasmic stores in brain endothelial cells *in vitro* to provide the local, high concentration necessary for recruitment of CXCR3+ leukocytes. This process also requires LFA-1 activity. In the absence of Caveolin-1, endothelial CXCL10 is secreted, and the local signaling cues are lost. Consistent with our *in vitro* data, genetic ablation of Caveolin-1 in endothelial cells reduces the severity of active experimental autoimmune encephalomyelitis (EAE), a murine model for multiple sclerosis, by decreasing the infiltration of CXCR3+ T cells into the CNS. Moreover, loss of Caveolin-1 protects against the adoptive transfer of autoreactive T cells. Our findings establish a novel mechanism by which brain endothelial cells utilize Caveolin-1 dependent CXCL10 intracellular stores to license T cells for transcellular migration across the blood-brain barrier.

## Introduction

Migration of T cells to sites of inflammation is a multistep process initiated by chemokine stimulation, ultimately resulting in diapedesis across the blood vessel endothelium (Mapunda et al., 2022). In the central nervous system (CNS), entry of proinflammatory CD4+ T cells can damage neurons and myelin, with adverse outcomes for neurological function (Mapunda et al., 2022). Adherent T cells crawl along the surface of the brain endothelium, extending podocytes and probing for sites favorable to diapedesis (Marchetti & Engelhardt, 2020; Martinelli et al., 2014). Subsequent leukocyte diapedesis occurs by transcellular migration (directly through the endothelial cytoplasm) or paracellular migration across cell-cell junctions (Liebner et al., 2018; Lutz et al., 2017). The crawling step and diapedesis step are differently governed by chemokine stimulation (Girbl et al., 2018). For example, luminal presentation promotes leukocyte adhesion (Sorensen et al., 2018), whereas subsequent exposure to a discrete chemokine pool influences diapedesis (Schoppmeyer et al., 2022; Shulman et al., 2011).

CXCL10 is an interferon-inducible chemokine that promotes activation and extravasation of CXCR3+ leukocytes to sites of tissue inflammation in autoimmune and infectious diseases. Perivascular hotspots of CXCL9 and CXCL10 promote CXCR3+ T cell migration by inducing inside-out activation of LFA1 and interaction with Intracellular Adhesion Molecule 1 (ICAM1) (Ngwenyama et al., 2019; Prizant et al., 2021). Endothelial cells themselves can also present chemokine depots to support integrin activation and T cell probing of the endothelium, creating a specific microanatomic location favorable for transmigration (Marchetti et al., 2022; Schoppmeyer et al., 2022; Shulman et al., 2011). Interestingly, chemokine presentation by activated endothelial cells promotes transcellular rather than paracellular diapedesis (Abadier et al., 2015; Marchetti et al., 2022), whereas new evidence suggests that homeostatic conditions with very low inflammatory chemokine production favor paracellular migration (Mapunda et al., 2022; Marchetti et al., 2022). There is an incomplete understanding of the mechanisms governing formation of intracellular chemokine stores and how these contribute to transcellular or paracellular BBB permeability.

In inflamed brain endothelial cells (BEC), transcellular cell migration and transcytosis of macromolecules are positively regulated by the integral membrane protein Caveolin-1 (Cav-1). Upon activation, Cav-1 undergoes a conformational change that triggers membrane curvature and formation of intracellular vesicles that contain cargos (Ohi & Kenworthy, 2022; Zimnicka et al., 2016). Suppression of brain endothelial cell Cav-1 is a key mechanism of BBB development, and loss of Cav-1 causes transcellular BBB permeability (Andreone et al., 2017; Chang et al., 2022; Chow & Gu, 2017; Wang et al., 2020). We and others have shown that Cav-1-related vesicular trafficking is normally maintained at low levels in mature brain EC but is upregulated in neuroinflammation *in vivo* (Knowland et al., 2014; Lutz et al., 2017; Salimi et al., 2020; Zhang et al., 2022).

It is unclear which steps of the leukocyte extravasation cascade are influenced by Cav-1. Caveolin complexes are enriched with ICAM-1 (Carman & Martinelli, 2015), and Cav-1 phosphorylation enhances leukocyte adhesion in an ICAM-1-dependent manner (Xu et al., 2013). In the experimental autoimmune encephalomyelitis (EAE) mouse model for multiple sclerosis, a disease caused by CNS infiltration of myelin-reactive leukocytes, mice with global deficiency in Cav-1 have less severe clinical disability, BBB leakage, and demyelination (Lutz et al., 2017; Wu et al., 2016). Interestingly, we found Cav-1 enhances autoreactive Th1 but not Th17 CD4+ T cell infiltration of the CNS (Lutz et al., 2017). Other groups have also identified Th1-specific features in CD4+ effector T cell motility and extravasation (Dusi et al., 2019; Gaylo-Moynihan et al., 2019). The observation that CXCR3 expression is high in Th1 CD4+ T cells (Rahimi & Luster, 2018) prompted us to hypothesize that intercellular interactions involving Cav-1 and CXCR3 could enhance transcellular diapedesis across the BBB.

In spite of these data, the specific role of endothelial cell Cav-1 in neuroinflammation is unknown because Cav-1 is expressed in many cell types. Cav-1 scaffolds costimulatory receptors at the immunological synapse to facilitate T cell activation (Borger et al., 2017; Schonle et al., 2016; Tomassian et al., 2011). Neuronal expression of Cav-1 protects against diabetes-associated cognitive dysfunction and amyloid β accumulation (Bonds et al., 2019; Head et al., 2010). We therefore created an endothelial cell specific conditional knockout of Cav-1 to assess the role of endothelial Cav-1 for T cell migration across the BBB.

In this study, we show that CXCL10 stimulation of CXCR3+ T cells enhances transcellular but not paracellular chemotaxis across BECs in a Cav-1 dependent manner. We find that endothelial Cav-1 is required for the expansion of intracellular CXCL10 stores in response to inflammation. Moreover, we find ICAM-1/LFA-1 interaction downstream of CXCR3 recruits T cells to transcellular migratory routes across BBB endothelia. Furthermore, our data establish that endothelial Cav-1, but not T cell Cav-1, dictates the severity of BBB permeability and neuroinflammation. Taken together, this work clarifies the mechanism by which CXCL10 and Cav-1 facilitate vascular inflammation and permeability of the BBB in association with neuroinflammatory diseases.

## Material & Methods

### Mice

All animal studies were approved by the UIC Animal Care and Use Committee. Endothelial-specific conditional Cav-1 knock-out (ECKO) mice were generated by crossing Cav-1^lox/lox^ mice (Cao et al., 2003) with Endo^SCL^CreERT2^+/-^ mice (Gothert et al., 2004) on a C57BL/6 background. Tamoxifen (Fisher MP215673891) (75mg/kg) was administered intraperitoneally every second day for 8 days beginning 28 days post-natal. Comparisons were made between Cav-1 ECKO mice (Endo^SCL^CreERT2^+/-^; Cav-1^lox/lox^) and control Cav-1 WT mice (Endo^SCL^CreERT2^-/-^; Cav-1^lox/lox^), all treated with Tamoxifen. Some experiments to measure Cre recombination used mice with genotype Endo^SCL^CreERT2^-/-^; Cav-1^lox/lox^; mTomato/mGFP^+/-^ (Jackson Laboratory 007576) (Muzumdar et al., 2007) in conjunction with immunofluorescent staining for Cav-1 protein. mTomato/mGFP mice are transgenic for genetically encoded Cre activity fluorescent reporter, in which membrane-anchored Tomato red fluorescent protein is transcribed after Cre-mediated deletion of an intervening enhanced GFP signal and stop codon flanked by LoxP sites. Wild-type C57Bl/6, Cav-1^-/-^, and CXCR3 KO (Jackson Laboratory 000664, 004585, 005796 respectively) mice were purchased from The Jackson Laboratory. Cav1^-/-^ mice were backcrossed 9 generations by outbreeding to C57Bl/6J.

### Active EAE

EAE was induced in 8-12 week old mice by subcutaneous injection with 100 μL of inoculant containing 100μg myelin oligodendrocyte glycoprotein peptide fragment 35-55 (MOG_35-55_, Thermo Scientific J66557MCR) in PBS with complete Freund’s adjuvant (CFA) containing 100μg *M. tuberculosis* H37Ra (BD 231141) (Lutz et al., 2012; Lutz et al., 2017). Mice received intraperitoneal injections of 100 uL 250 ng/μL toxin from *B. pertussis* (List Biological Laboratories 181) at 0- and 2-days post-immunization (DPI). MOG/CFA and pertussis were administered between 3:00 pm and 6:00 pm. Male and female mice were used. Mice were examined for clinical signs of EAE using a 0-5 scale with 0.5 point gradations. 0: no signs, 1: flaccid tail, 2: hind limb paresis, 3: hind limb paralysis, 4: hind and forelimb paralysis, 5: moribund.

### Passive EAE

Cells from spleen and lymph nodes (inguinal, cervical, axillary) were collected from 8-12 week old C57BL/6 mice 7 days after immunization with MOG_35-55_ in CFA without *B. pertussis* toxin. Cells from sex-matched donor mice were cultured in RPMI with 5% FBS, L-glutamine, beta-mercaptoethanol, non-essential amino acids, 20 μg/mL MOG_35-55_, 1 ng/mL IL-12 (Biolegend 577002), and 5 ng/mL IL-2 (Biolegend 575404) at 1×10^7^ cells/mL for 72 hrs. Lymphoblasts were counted based on cell size and morphology. Cells were resuspended in PBS at 1×10^8^ lymphoblasts/mL. Sex-matched 8–12-week-old C57BL/6 or Cav-1^-/-^ recipient mice received a single 200 μL administration of 2×10^7^ T cell blasts by retro-orbital injection. Recipient mice also received intraperitoneal injections of 100 μL (250 ng/μL) toxin from *B. pertussis* at 0 and 2 DPI.

### Flow cytometry

Brains and spinals cords were dissected and mechanically homogenized between ground glass slides. Mononuclear cells were isolated at the interphase of a 30%–70% Percoll gradient (GE Healthcare P1644). Splenocytes from were isolated and homogenized through a 70 μm cell strainer. Red blood cells were lysed with ammonium-chloride-potassium buffer. Single cell suspensions from brain, spinal cord, and spleen were prepared for flow cytometry. Fixable viability dye (Zombie Green, Biolegend 423111) and Fc receptor blockade (anti-mouse CD16/32, Biolegend 101319) were followed by incubation with antibodies against CD45 (Biolegend 103154), CD4 (BD Horizon 563151), CXCR3 (Biolegend 126521), CXCR4 (Biolegend 146505), CCR6 (Biolegend 129815), CCR2 (Biolegend 150627), CD11b (Biolegend 101229). Fixation and permeabilization (Biolegend 421403) was followed with an antibody against Cav-1 (Cell Signaling 31411S). Cells were analyzed with a Beckman CytoFLEX S flow cytometer.

### Primary cell culture

For T cell cultures, splenocytes from C57BL/6 or CXCR3 KO mice were collected 7 days after immunization with MOG_35-55_ in CFA without *B. pertussis* toxin. Splenocytes were cultured in RPMI with 5% FBS, L-glutamine, beta-mercaptoethanol, non-essential amino acids, 20 μg/mL MOG_35-55_, and 1 ng/mL IL-12 (Biolegend 577002) for 3 days to favor Th1 differentiation. 5 ng/mL IL-2 (Biolegend 575404) was added to media and cells were cultured another 2 days (Lutz et al., 2017). T cell cultures were enriched to ∼50% CXCR3+ CD4+ T cells after 5 days *in vitro* (Fig. S3 A-D).

Primary mouse brain endothelial cells (mBECs) were isolated as described in detail (Marottoli et al., 2021). Briefly, cerebral cortices were isolated from 3-5 week old C57Bl/6 or Cav-1^-/-^ mice, minced, and digested with papain (Worthington Biochemical LK003178) and DNase (Worthington Biochemical LK003172). Digested tissue was homogenized, microvessels were isolated by 25% BSA gradient centrifugation, and red blood cells were lysed with ammonium-chloride-potassium buffer. Microvascular endothelial cells were cultured on glass 12-chamber slides (Ibidi 81201) coated with poly-D-lysine (Sigma P7280), fibronectin (Sigma F0895), collagen-I (Sigma C8919), and laminin (L2020) in complete endothelial cell media (Lonza CC-4147, Lonza CC-3156). Inclusion of puromycin (10 μg/mL) (Sigma P8833) during the first 24 hours of culture eliminated non-endothelial cells. Tumor necrosis factor alpha (TNFα) 1 ng/mL (Biolegend 575206) was added to mBECs 16 hours before T cell migration experiments (Lutz et al., 2017; Martinelli et al., 2014).

### ELISA

Blood was collected from WT and Cav-1^-/-^ mice with EAE at 0, 5, and 10 DPI by tail snips in tubes with 10 μL 0.5 mM EDTA, and plasma was prepared by centrifugation at 2000xg for 15 min. Primary WT and Cav-1^-/-^ mBECs were isolated and cultured as detailed above. After 5 days in culture, IFNγ (1000 U/mL) (Biolegend 575302) was added to cells in complete endothelial cell media. After 24 hr of IFNγ treatment, cell supernatant was collected for ELISA and mBECs were fixed and stained for immunofluorescence microscopy analysis. CXCL10 was measured in plasma and cell supernatant using a DuoSet ELISA Kit (R&D Systems DY466-05, R&D Systems DY008B). ELISAs were performed according to manufacturer’s protocol.

### Static transmigration

Primary CD4+ T cell cultures from C57BL/6 or CXCR3 KO mice were incubated with mBECs from C57BL/6 or Cav-1^-/-^ mice. 50,000 CD4+ T cells were added to each well of mBECs in 12-well chamber slides in Hank’s balanced salt solution (HBSS) with 1% FBS supplemented with 5 nM CXCL10 (Biolegend 573604) or vehicle (PBS). In some experiments, mBECs were pretreated with 5 nM methyl-ß-cyclodextrin (Sigma C4555) for 1 hr prior to incubation with CD4+ T cells. In some experiments, CD4+ T cells were pretreated with anti-CD18 blocking antibody (Biolegend 101418) for 20 min prior to incubation with mBECs. CD4+ T cells were allowed to adhere and transmigrate for 60min at 37°C. Cultures were washed with HBSS three times and fixed with 4% PFA before immunostaining. Colorimetric development was detected with a Beckman Coulter DTX880 plate reader. CXCL10 concentration was normalized to total protein concentration in each sample as assessed with Pierce BCA Protein Assay Kit (Thermo 23227).

### Immunofluorescence imaging

Co-incubated primary CD4+ T and mBECs were fixed with 4% paraformaldehyde in PBS for 15 minutes before staining for immunofluorescence imaging. Fixed cultures were stained with antibodies against CD45 (1:500 Millipore Sigma 05-1416), Zonula occludens-1 (ZO-1, 1:250 Invitrogen 33-9100), and Cav-1 (1:500 Invitrogen PA5-17447). Images were acquired with a Leica DMI8 microscope and 40X Z-stack images were processed with Leica LIGHTNING deconvolution software. Processed images were analyzed for T cell paracellular and transcellular migration across mBECs using Leica LASX and Fiji software.

For CXCL10 immunostaining, mBECs treated with IFNγ were fixed with 4% paraformaldehyde in PBS for 15 minutes and stained with antibodies against Zona occludens-1 (ZO-1, 1:250 Invitrogen 33-9100) and CXCL10 (1:500 Abcam ab9938). Images were acquired with a Zeiss LSM880 confocal microscope with Airyscan enhanced resolution. Four regions were selected from each image for CXCL10 analysis. For assessment of cytoplasmic CXCL10 depots, perinuclear endoplasmic reticulum and Golgi regions were excluded. CXCL10 particle size in selected regions was analyzed with Fiji software.

### Microfluidic transmigration

Primary mBECs were cultured in Ibidi 6-channel slides (Ibidi 80606) coated with collagen-I (Sigma C8919) for 4 days with complete EGM-2 media (Lonza CC-4147, Lonza CC-3156). Upon reaching 80-90% confluence, media was changed to complete EGM-2 with 1 ng/mL TNFα (Biolegend 575206) and shear stress of 1 dyn/cm^2^ was incorporated using a microfluidic pump (Ibidi 10902) for 24h. Primary CD4+ T cells stained with fixable MemBrite 594/615 (Biotium 30096) were injected into the microfluidic circulation in HBSS with 1% FBS supplemented with 5 nM CXCL10 (Biolegend 573604) or vehicle (PBS). In some experiments, CD4+ T cells were pretreated with anti-CD18 blocking antibody (Biolegend 101418) for 20 min. Microfluidic flow was paused for 1 minute to allow CD4+ T cells to adhere before shear stress of 1 dyn/cm^2^ resumed. Transmigration was permitted under flow conditions for 60 min at 37°C with 5% CO2. Several regions in each channel were selected and imaged once per minute with a Leica DMI8 microscope.

### IMARIS analysis

IMARIS Microscopy Analysis Software (Oxford Instruments) was used for analysis and automated cell tracking of live *in vitro* transmigration. Fluorescent T cell movement was tracked using IMARIS spot-detection function in timelapse images. Automated cell tracking with autoregressive motion was manually edited to remove redundancies and fill gaps in tracks resulting from insufficient fluorescence detection. Meandering index was calculated as total track displacement/total track length. Instantaneous velocity was measured by distance traveled between successive timepoints. For cell arrest calculations, cells moving at a speed less than 0.1 μm/s were considered arrested.

## Results

### CXCL10 enhances adhesion and transcellular migration of CD4+ T cells across mBECs

T cell migration through the BBB involves sequential steps of rolling, adhesion and crawling on the endothelial surface, extension of membrane protrusions into the endothelium, and diapedesis either through (transcellular) or between (paracellular) endothelial cells (Mapunda et al., 2022). Endothelial intrinsic factors can instruct and guide T cell route of transmigration (Charabati et al., 2022; Marchetti et al., 2022; Schoppmeyer et al., 2022). However, less is known about how T cell signaling directs paracellular and transcellular diapedesis decisions. We previously showed that activated CD4+ Th1 T cells, known to highly express the chemokine receptor CXCR3, favor Cav-1 dependent transcellular migration (Lutz et al., 2017). We investigated if CXCR3 signaling directs transcellular diapedeses across BECs. Primary MOG_35-55_-specific CD4+ T cells were cultured from WT or CXCR3 deficient (CXCR3 KO) mice and differentiated into Th1-like cells by culturing with IL-12, IL-2, and MOG_35-55_ for 5 days. WT and CXCR3 KO CD4+ Th1 T cells were applied to a monolayer of primary mBECs in the presence of 5 nM CXCL10. After 1-hour, non-adherent cells were removed and adherent CD4+ T cells were quantified by immunostaining for CD45 (Fig. 1A). As expected, adhesion of WT but not CXCR3 KO CD4+ T cells to mBECs was enhanced with CXCL10 (Fig. 1B). Z-stacks were acquired to analyze the diapedesis route (Fig. 1A). CD4+ T cells with >50% of their membrane protrusions colocalizing with junctional protein ZO-1 were categorized as paracellular migration events. CD4+ T cells were considered transcellular if they were embedded in the mBEC monolayer with less than 10% of their protrusion surface colocalizing with ZO-1. Transcellularly migrating cells were frequently flanked by endothelial Cav-1 (Fig. 1A). We found that CXCL10 stimulation significantly increased transcellular CD4+ T cell interactions with mBECs and that this increase was abolished in CXCR3 KO CD4+ T cells (Fig. 1C). These data suggest that CXCR3 activation by CXCL10 specifically primes CD4+ T cells for transcellular migration across the BBB.

**Figure 1:**
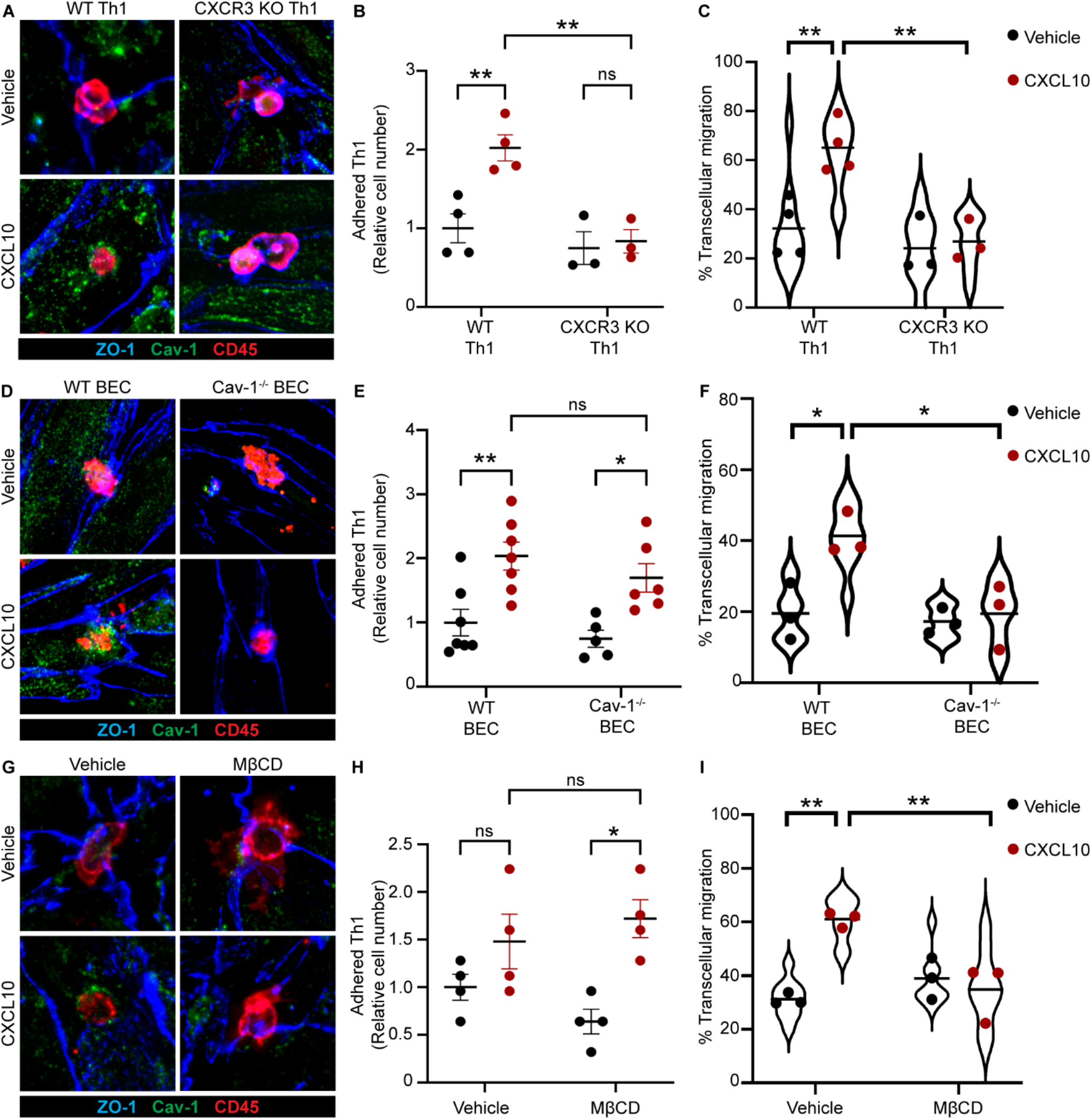
CXCL10-mediated transcellular migration of CD4+ T cells is dependent on endothelial Cav-1. **(A)** Representative immunofluorescent images of WT and CXCR3 KO Th1 CD4+ T cell transmigration across mBECs in the presence of 5 nM CXCL10 or vehicle. Immunostaining was conducted for ZO-1 (blue), Cav-1 (green), and CD45 (red) (A, D, G). Examples show T cells engaging with ZO-1+ paracellular junctions in the absence of CXCL10, and T cells engaging with Cav-1+ transcellular sites in the presence of CXCL10. **(B)** Adhesion of WT and CXCR3 KO CD4+ T cells to mBEC, with 5 nM CXCL10 or vehicle. Points represent averages from 3-4 independent experiments. Adherent cell count includes migratory CD4+ T cells. **(C)** Relative frequency of transcellular migration of WT and CXCR3 KO CD4+ T cells stimulated with 5 nM CXCL10 or vehicle. Points represent averages from 3-4 independent experiments; violin plots depict distribution of all data from all experiments. **(D)** Representative images of CD4+ T cell transmigration across WT and Cav-1^-/-^ mBECs with 5 nM CXCL10 or vehicle. **(E)** Quantitation of CD4+ T cells adherent to WT and Cav-1^-/-^ mBECs in the presence of 5 nM CXCL10 or vehicle. Points represent averages from 4-7 independent experiments. Adherent cell count includes migratory CD4+ T cells. **(F)** Relative frequency of transcellular migration of CD4+ T cells across WT and Cav-1^-/-^ mBECs in the presence of 5 nM CXCL10 or vehicle. Points represent averages from 3 independent experiments, violin plot represents distribution of all data from all experiments. **(G)** Representative images of CD4+ T cell transmigration across WT and MβCD-treated mBECs in the presence of 5 nM CXCL10 or vehicle. **(H)** Quantitation of CD4+ T cell adhesion to WT and MβCD-treated mBEC with 5 nM CXCL10 or vehicle. Points represent averages from 4 independent experiments. Adherent cell count includes migratory CD4+ T cells. **(I)** Relative frequency of transcellular migration of CD4+ T cells across WT and MβCD-treated mBECs with 5 nM CXCL10 or vehicle. Points represent averages from 3 independent experiments; violin plot represents all individual data points from all experiments. One-way ANOVA with Sidak’s post-hoc comparisons between groups. * p<0.05 ** p<0.01

### CXCL10-mediated transcellular migration of CD4+ T cells is dependent on mBEC Cav-1

Publications from our group and others have shown the importance of Cav-1 in BBB permeability (Andreone et al., 2017; Guerit et al., 2021; Lutz et al., 2017). However, the mechanism by which Cav-1 regulates neuroinflammation or BBB transendothelial migration is unclear. To interrogate the role of Cav-1 in transcellular migration of CXCR3+ CD4+ T cells across BBB ECs, we stimulated MOG_35-55_-specific CD4+ Th1 T cells with 5 nM CXCL10 immediately prior to application to WT or Cav-1^-/-^ mBECs. We again analyzed transcellular and paracellular interactions of CD4+ T cells with BEC with immunostaining and microscopy (Fig. 1D). Endothelial Cav-1 deficiency had no effect on CXCL10-mediated CD4+ T cell adhesion to mBECs (Fig. 1E). We found that transcellular migration of CXCL10-stimulated CD4+ T cells was inhibited in Cav-1^-/-^ mBECs (Fig. 1F). This indicates that CXCL10 stimulation of transcellular migration of CD4+ T cells across the BBB requires endothelial cell Cav-1 expression.

Cav-1 is a membrane anchored protein that forms complexes which regulate signal transduction and endocytic vesicle trafficking (Ohi & Kenworthy, 2022). These complexes, called caveolae, form within cholesterol-enriched membrane microdomains in cells expressing Cav-1. Cyclodextrins are cyclic oligosaccharides commonly used as chelating agents. Methyl-β-cyclodextrin (MβCD) binds to cholesterol and free fatty acids in plasma membranes and has been widely used as a pharmacologic approach to disrupt caveolae (Cassuto et al., 2014; Feher et al., 2010). Here, we utilized MβCD to begin to investigate the function of mBEC Cav-1 expression in transcellular migration of CD4+ Th1 T cells. mBECs were treated with 10 mM MβCD for 1 hour prior to the introduction of MOG_35-55_-specific CD4+ T cells that had been stimulated with 5nM CXCL10. After 1 hour, adherent CD4+ T cells were quantified and analyzed for paracellular and transcellular migration by immunofluorescence microscopy (Fig. 1G). We found that MβCD did not inhibit CXCL10-stimulated adhesion of CD4+ T cells to mBECs (Fig. 1H), in agreement with data obtained in our targeted genetic approach (Fig. 1E) further suggesting that endothelial caveolae are not required for or regulate adhesion of chemokine-activated CD4+ T cells. However, MβCD disruption of caveolae in mBECs inhibited transcellular diapedesis of CD4+ T cells in the presence of CXCL10 (Fig. 1I). Together, data generated using genetic and pharmacologic approaches to disrupt endothelial Cav-1 expression or functional caveolae suggests that Cav-1 plays a role in regulating transcellular migration of chemokine-activated CD4+ T cells following firm adhesion to mBEC. How Cav-1 directs adherent CD4+ T cells transmigration, however, is still unknown.

### CD4+ T cell transcellular migration across mBECs is dependent on LFA-1/ICAM-1 signaling

In leukocytes, chemokine receptor signaling leads to inside-out activation of integrins. For CXCR3, stimulation with CXCL10 results in activation LFA-1, an important integrin in T cell migration. The cognate receptors for LFA-1 on ECs are ICAM-1 and ICAM-2. Interaction between LFA-1 and ICAM-1 facilitates firm adhesion to endothelia (Gerard et al., 2021). Here, we investigate if CD4+ Th1 T cell LFA-1 interaction with endothelial cell ICAM is necessary for CXCL10-mediated transcellular BEC migration. MOG_35-55_-specific CD4+ T cells were treated with 40 ng/mL integrin β_2_ blocking antibody (anti-CD18) to inhibit LFA1 prior to addition of T cells to mBECs, in the presence of 5 nM CXCL10. CD4+ T cell adhesion and transmigration across the mBEC monolayer was analyzed by immunofluorescence imaging (Fig. 2A). Anti-CD18 did not eliminate adhesion, suggesting additional and alternative interactions including selectins, VCAM, and antigen presentation also contribute to initial firm adhesion (Fig. 2B) (Heng et al., 2022; Manes et al., 2022; Mapunda et al., 2022). We went on to assess the role of LFA-1 on transcellular and paracellular diapedesis. We found that blocking LFA-1 with anti-CD18 impeded transcellular migration of CXCL10-activated MOG_35-55_-specific CD4+ T cells across mBECs (Fig. 2C). Together with previous data, this suggests CXCL10/CXCR3 T cell activation promotes transcellular, Cav-1 dependent diapedesis through BBB BEC by a mechanism involving LFA-1/ICAM.

**Figure 2:**
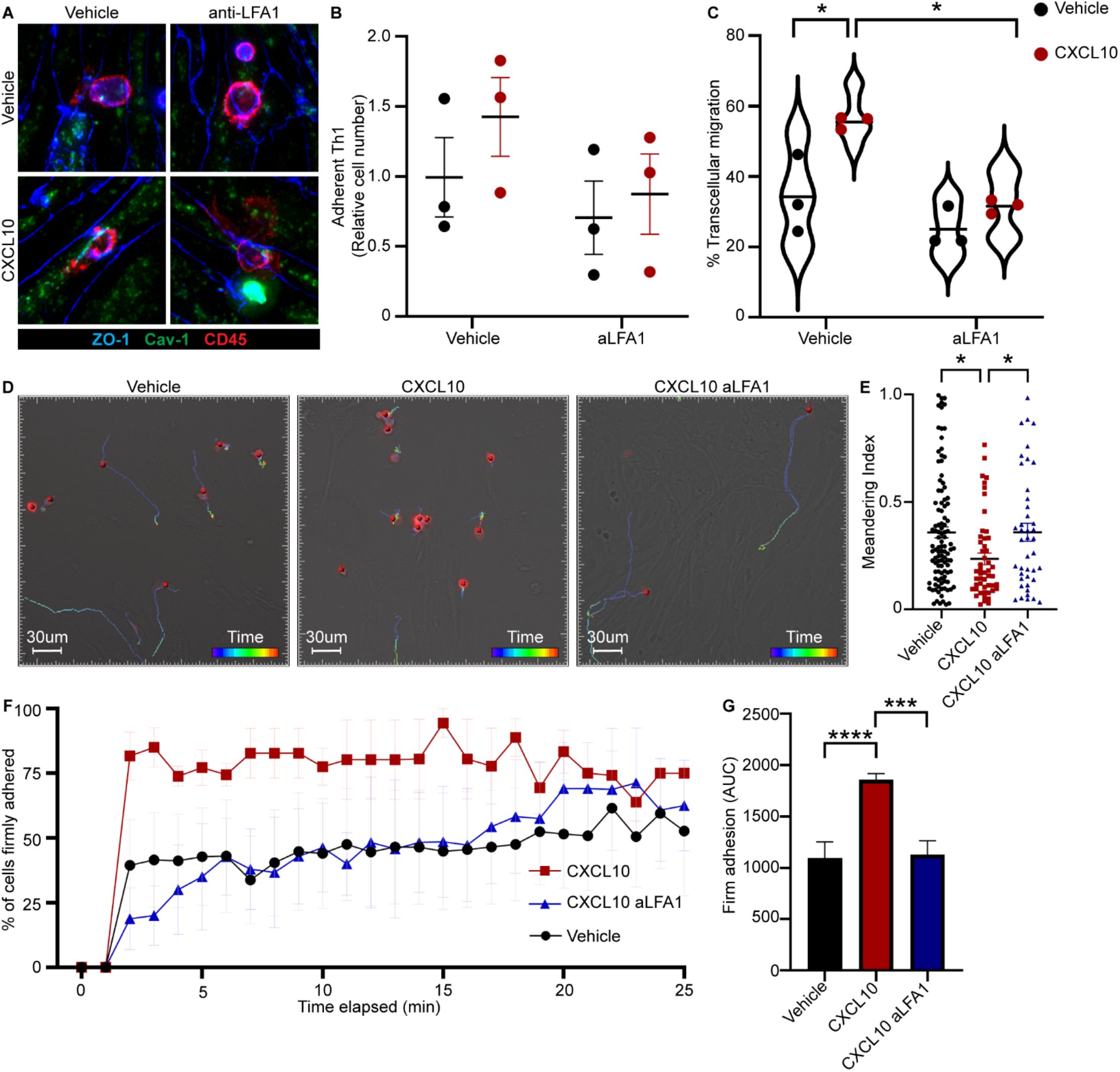
LFA-1 ICAM-1 interaction facilitates CXCL10-mediated CD4+ T cell transcellular migration. **(A)** Representative images of vehicle- or anti-LFA-1 treated CD4+ T cells transmigrating across mBECs in the presence of 10 nM CXCL10 or vehicle. Immunostaining was conducted for junctional ZO-1 (blue), Cav-1 (green), and CD45 (red). **(B)** Adhesion of vehicle- or anti-LFA-1 treated CD4+ T cells to mBEC in the presence of 5 nM CXCL10 or vehicle. Points represent the average from n=3 experiments. Adherent cell count includes migratory CD4+ T cells. **(C)** Relative frequency of transcellular migration of vehicle- or anti-LFA-1 treated CD4+ T cells across mBECs in the presence of 5nM CXCL10 or vehicle. Points represent averages from n=3 experiments; violin plot represents the distribution of all data from all experiments. **(D)** Representative IMARIS track plots from microfluidic transmigration of vehicle, CXCL10, and anti-LFA-1 treated CD4+ T cells incubated with mBECs. (**E)** Quantitative analysis of meandering index (total track displacement/total track length) of vehicle, CXCL10, and anti-LFA-treated CD4+ T cells. **(F)** Analysis of time to firm adhesion of vehicle, CXCL10, and anti-LFA-1 treated CD4+ T cells incubated with mBECs in the presence of 5nM CXCL10 or vehicle. **(G)** Area under the curve analysis of firmly adhered CD4+ T cells from 0-25 minutes. One-way ANOVA with Sidak’s post-hoc comparisons between groups. * p<0.05 *** p<0.001 **** p<0.0001

ICAM-1 clustering and crosslinking on endothelial cells has been shown to increase Cav-1-dependent transcytosis and signaling, resulting in increased endothelial permeability (Hu et al., 2008). Our data suggest that specific integrin-CAM interactions, which are often required for firm adhesion, may also direct the route of leukocyte migration across endothelia. We sought to clarify the role of LFA-1 in transendothelial migration of CD4+ T cells by imaging transmigration in microfluidic chambers with physiological shear forces. Primary mBECs were cultured in Ibidi channel slides with shear stress resembling post-capillary venule flow for 24 hours prior to and during transmigration of primary MOG_35-55_-specific CD4+ Th1 T cells. CD4+ T cells were pretreated with 40 ng/mL anti-CD18 blocking antibody (or vehicle) and applied to mBECs with or without 5 nM CXCL10. After an initial pause in flow to promote adhesion, chambers were imaged at 1 frame/minute for 1 hour under continuous flow. CD4+ T cell crawling duration, speed, and direction were analyzed using IMARIS imaging analysis software (Fig. 2D). We evaluated the meandering index (track length (μm)/track displacement (μm)) as a measure of purposeful directionality of CD4+ T cell crawling. We found that CD4+ T cells treated with CXCL10 had lower meandering than untreated or anti-CD18 treated CD4+ T cells (Fig. 2E), suggesting LFA-1 interaction with BECs directs crawling of CXCL10-activated CD4+ T cells. We next interrogated the duration of CD4+ T cell crawling to measure the elapsed time prior to transendothelial migration. At each timepoint, the speed of the cell (μm/s) was measured by displacement from the previous frame. Stationary, adherent cells (<0.1 μm/s) were categorized as arrested; the few cells that detached at this time were counted separately. We found that untreated and anti-CD18 treated CD4+ T cells took significantly longer to arrest than CXCL10-treated CD4+ T cells (Fig. 2F-G). This result suggested that LFA-1 influences how CD4+ T cells search for transmigration locations through BECs. LFA-1 activation downstream of CXCR3 signaling primed CD4+ T cells for Cav-1-dependent transcellular migration across mBECs. Without LFA-1/ICAM-1 interaction, adherent CD4+ T cells exhibited prolonged crawling without diapedesis (Martinelli et al., 2014).

### BEC Cav-1 organizes intracellular CXCL10 depots

Chemokines including CXCL10 are upregulated in the cerebrovasculature in neuroinflammatory disease (Heng et al., 2022; Karin, 2020; Liebner et al., 2018; Manes et al., 2022). The spatial distribution of CXCL10 influences whether infiltrating leukocytes can cross the brain EC, are retained in perivascular spaces, or are able to enter the brain parenchyma (Giladi et al., 2020; Heng et al., 2022; Muller et al., 2010). Endothelial cells outside of the nervous system are known to promote transendothelial T cell migration by the regulated release of intracellular chemokine stores at locations of intimate leukocyte-endothelial interaction (Schoppmeyer et al., 2022; Shulman et al., 2011). Whether endothelial cell Cav-1 plays a role in chemokine presentation or secretion from the BBB is unknown. Strikingly, it was recently reported that dephosphorylation of Cav-1 enhances CXCL10 secretion from mesenchymal stem cells (Wang et al., 2021). We therefore investigated if reduction in Cav-1 expression altered CXCL10 aggregation or secretion in brain endothelial cells.

CXCL10 distribution was analyzed in primary WT and Cav-1^-/-^ mBECs treated with interferon gamma (IFNγ), the canonical activator for CXCL10 production (Koper et al., 2018). Abundance and size of cytoplasmic CXCL10 “depots” (Schoppmeyer et al., 2022; Shulman et al., 2011) were interrogated in confocal Z-stack slices (Fig. 3A). In WT mBECs, IFNγ stimulation increased the diameter of CXCL10 depots (Fig. 3B) and total CXCL10 immunopositive area (Fig. 3C). In contrast, there was no expansion of CXCL10 depot diameter in IFNγ-treated Cav-1^-/-^ mBECs (Fig. 3B), nor was there an increase in area of CXCL10 immunoreactivity (Fig. 3C). These data suggest that Cav-1 is required for brain endothelial cells to dynamically expand chemokine depots in response to inflammation.

**Figure 3:**
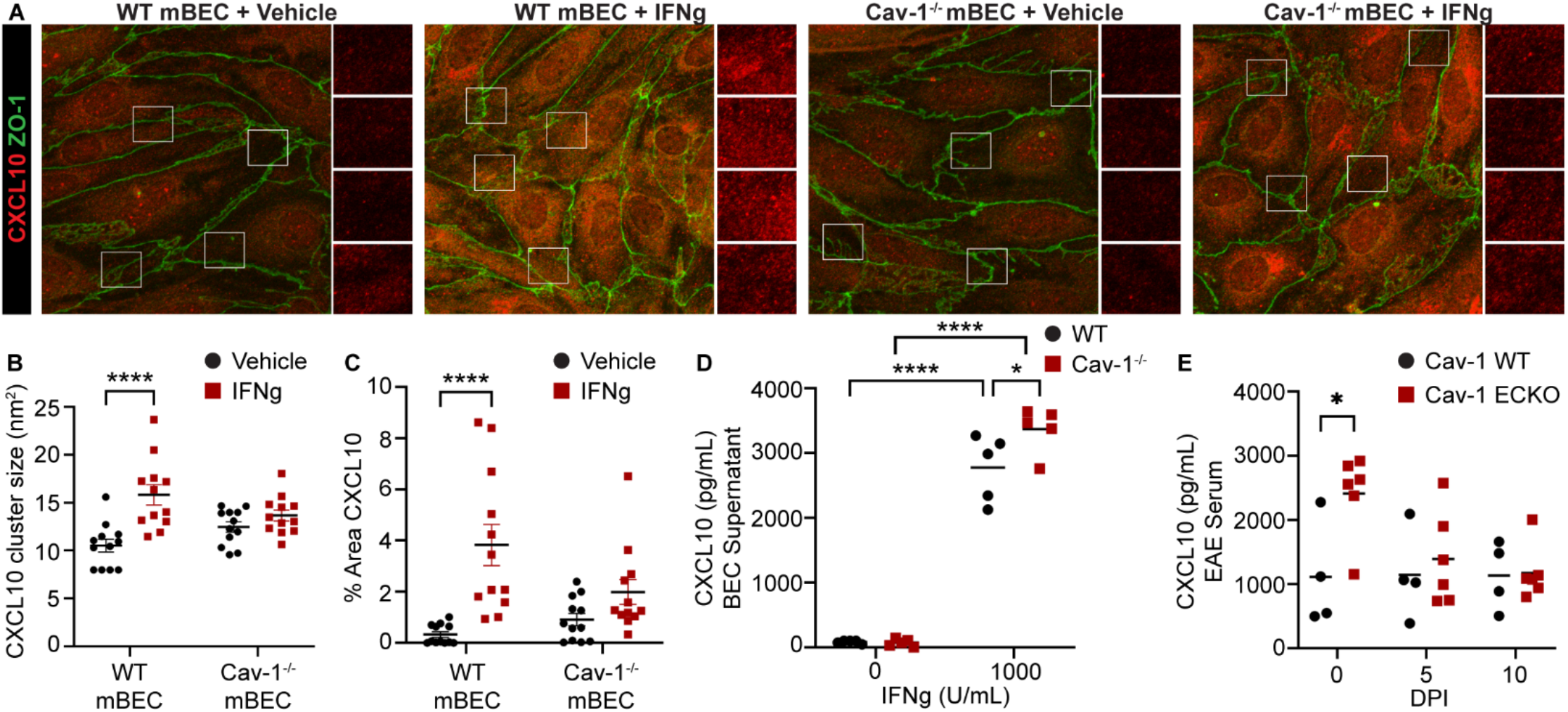
Endothelial Cav-1 functions in CXCL10 cytoplasmic aggregation and presentation. **(A)** Representative images of WT and Cav-1^-/-^ mBECs treated with IFNγ or vehicle and immunostained for CXCL10 (red) and ZO-1 (green). White squares are regions used for CXCL10 analysis, shown magnified to the right of main image. **(B)** CXCL10 cluster size (nm^2^) in IFNγ-treated WT and Cav-1^-/-^ mBECs. Two-way ANOVA with Sidak’s post-hoc comparisons between WT and Cav1^-/-^. **(C)** CXCL10 percent area in white squares of IFNγ-treated WT and Cav-1^-/-^ mBECs. Two-way ANOVA with Sidak’s post-hoc comparisons between WT and Cav1^-/-^. **(D)** CXCL10 ELISA of IFNγ-treated primary MBECs from WT and Cav-1^-/-^ mice. Supernatant collected 24hr after addition of IFNγ. One-way ANOVA with Sidak’s post-hoc comparisons. **(E)** ELISA of plasma from Cav-1 WT and Cav-1 ECKO mice at 0, 5, and 10 DPI. * p<0.05 **** p<0.0001

We reasoned that constitutive chemokine secretion would deplete intracellular stores yielding small chemokine depots, like we observed in Cav-1^-/-^ mBECs. We therefore measured whether CXCL10 secretion was regulated by Cav-1 expression. Supernatant from WT and Cav-1^-/-^ mBECs treated with IFNγ was collected for ELISA analysis of CXCL10. We found significantly more CXCL10 in the supernatant of IFNγ treated Cav-1^-/-^ mBECs as compared with WT mBECs (Fig. 3D).

CXCL10 levels within the blood are reported to be elevated in relapsing remitting MS patients in some but not all studies (Franciotta et al., 2001; Ghafouri-Fard et al., 2021). We analyzed CXCL10 levels in the blood of WT and Cav-1 ECKO mice during the onset of EAE. In our study, CXCL10 levels in the blood remained constant at 0, 5, or 10 days post-immunization (DPI) in WT mice with EAE (Fig. 3E). CXCL10 was increased in the blood of Cav-1 ECKO mice prior to immunization, but not during EAE, suggesting that chemokine levels in the blood are regulated during disease (Fig. 3E).

Together, these data suggest that Cav-1 promotes the organization of CXCL10 into cytoplasmic stores for efficient, local activation of CXCR3+ leukocytes. Loss of Cav-1 prevents CXCL10 depots from expanding and instead promotes secretion into the extracellular space. Stimulated, local release of intraendothelial chemokine stores plays an important role in transcellular T cell migration (Schoppmeyer et al., 2022). Therefore, reduced CXCL10 release during T cell interaction at the Cav-1 deficient BBB may preclude CXCR3+ CD4+ T cells from entering the CNS in the EAE mouse model of neuroinflammation.

### Endothelial Cav-1 contributes to neuroinflammation in EAE

As we and others have previously reported, Cav-1^-/-^ mice presented with lessened disease severity in the experimental autoimmune encephalomyelitis (EAE) model for multiple sclerosis induced by active immunization with MOG_35-55_ peptide (Lutz et al., 2017; Wu et al., 2016) (Fig. 4A-B). However, these studies could not establish the specific function of brain endothelial cell Cav-1 for neuroinflammation, because Cav-1 plays important roles in diverse cell populations including endothelial cells, T cells, B cells, and neurons (Ohi & Kenworthy, 2022).

**Figure 4:**
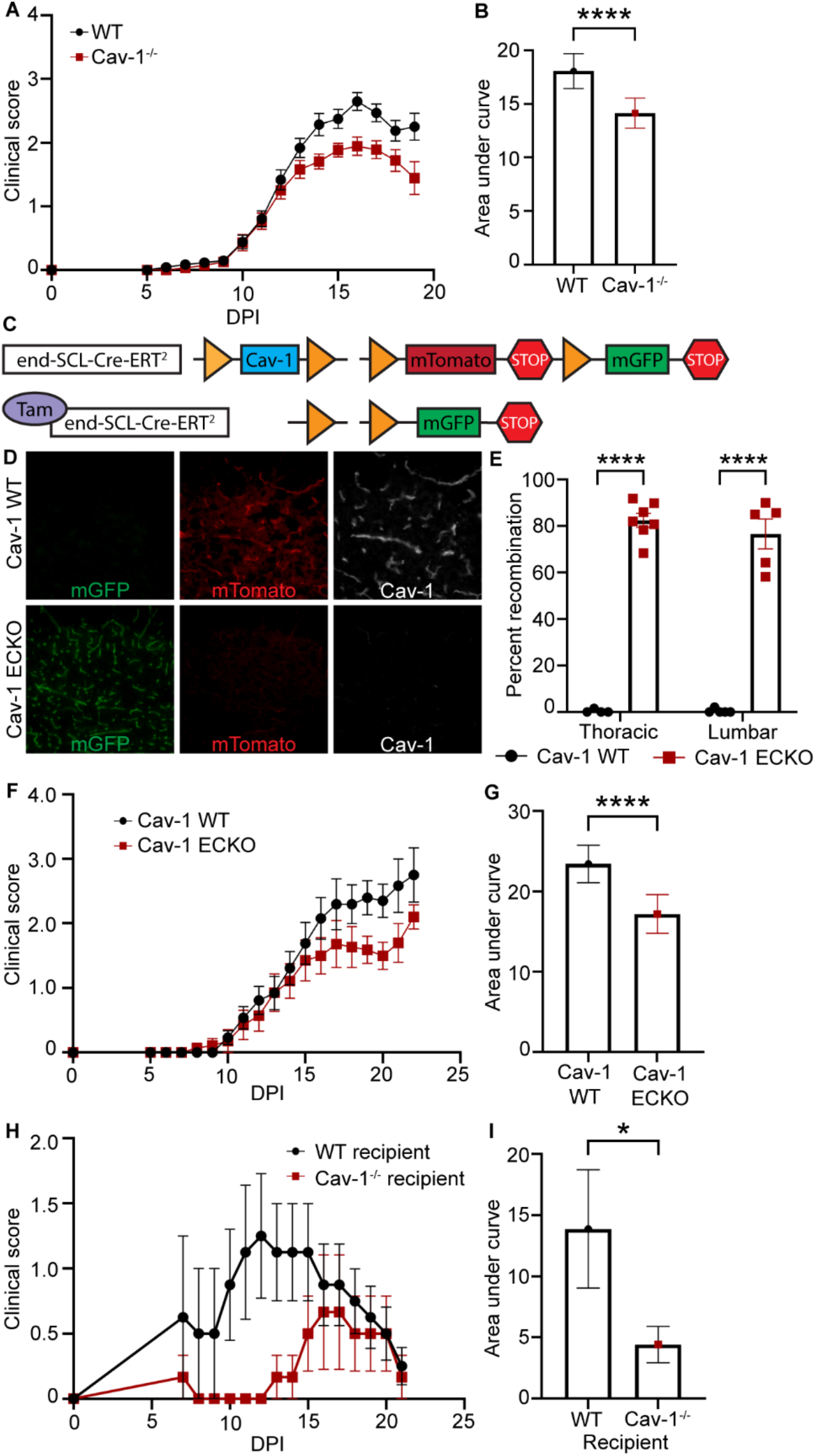
Endothelial Cav-1 contributes to neuroinflammation in EAE. **(A)** Clinical curve of EAE in WT (n=34) and Cav-1^-/-^ (n=29) mice. (**B)** Area under the curve analysis of WT and Cav-1^-/-^ clinical curves for 0-19 DPI. **(C)** Schematic of Cav-1 endothelial cell conditional knock-out (ECKO) genetic constructs. **(D)** Spinal cord section of tamoxifen-treated Cav-1 WT and Cav-1 ECKO mice. Immunofluorescent staining for Cav-1 (white) is shown with mGFP+ (green) recombined and mTomato+ (red) unrecombined vessels. **(E)** Quantification of mGFP to mTomato recombination in spinal cord blood vessels of tamoxifen-treated Cav-1 WT and Cav-1 ECKO mice. **(F)** Clinical curve of EAE in tamoxifen-treated Cav-1 WT (n=13) and Cav-1 ECKO (n=14) mice. **(G)** Area under the curve analysis of Cav-1 WT and Cav-1 ECKO clinical curves for 0-22 DPI. (**H)** Adoptive transfer EAE clinical curves of WT MOG-specific CD4+ T cells to WT (n=4) and Cav-1^-/-^ (n=3) recipient mice. (**I)** Area under the curve analysis of adoptive transfer EAE clinical curves 0-21 DPI. Unpaired t-tests (B, G, I) and one-way ANOVA with Sidak’s post-hoc comparison test (E). * p<0.05, **** p<0.0001

We therefore designed studies to determine whether the expression of Cav-1 in brain endothelial cells promotes disease severity by influencing leukocyte extravasation *in vivo*. For this, we established a Cav-1 endothelial-specific conditional knockout (Cav-1 ECKO) mouse strain (Fig. 4C). We crossed mice with LoxP sites flanking the Cav-1 locus (Cao et al., 2003), mice with the mTomato/mGFP Cre reporter (Muzumdar et al., 2007), and mice with endothelial-specific, inducible Cre (Gothert et al., 2004). Tamoxifen treatment of adult Cav-1 ECKO mice resulted in Cre-mediated deletion of Cav-1, and concordant switch from mTomato red fluorescence to mGFP green fluorescence, in CNS blood vessels (Fig. 4D-E). We found that active EAE in Cav-1 ECKO mice recapitulates the diminished progression and reduced severity of disease in Cav-1^-/-^ mice (Fig. 4F-G). Time to onset of disease signs was not significantly different between Cav-1 WT and Cav-1 ECKO mice (Fig. S2H). We found no difference in EAE disease progression or severity based on biological sex (Fig. S2D-G, S2I-L).

We also tested a passive model of EAE in which primary MOG_35-55_-specific T cells are cultured in Th1-polarizing conditions and adoptively transferred into healthy WT or Cav-1^-/-^ mice. We found that Cav-1^-/-^ mice that received MOG_35-55_-specific CD4+ T cells showed delayed onset of disease signs and reduced severity compared to WT recipients (Fig. 4H-I). Together, these data suggest that endothelial Cav-1 enhances neuroinflammation in EAE. Next, we investigated if endothelial Cav-1 regulates BBB permeability to encephalitogenic T cells in EAE.

### Cav-1 deficient mice have reduced CXCR3+ T cell infiltration in the CNS during EAE

We have previously reported that IFNγ+ CD4+ Th1 T cells have reduced transcellular BBB transmigration *in vitro* and entry into the CNS of Cav-1^-/-^ mice *in vivo* during EAE (Lutz et al., 2017). A distinguishing characteristic of Th1 cells is high expression of CXCR3 (Duckworth et al., 2021; Heng et al., 2022; Koper et al., 2018). Chemokines and their receptors spatially and temporally regulate migration of immune cells to sites of inflammation (Heng et al., 2022; Karin, 2020). To determine if T cells expressing specific chemokine receptors are excluded from the CNS in Cav-1^-/-^ mice during EAE, we examined the role of chemokine receptor expression on CNS-infiltrating and peripheral T cells throughout disease development.

In both WT and Cav-1^-/-^ mice, onset of disease signs in EAE occurred, on average, at 10 days post-immunization (DPI) (Fig. S2C). We first interrogated the role of Cav-1 in CNS-infiltration of CD4+ T cells during EAE onset and found that overall CD4+ T cells were reduced in both the brain and spinal cord of Cav-1^-/-^ mice (Fig. 5A-B). Further characterization revealed that CXCR3 expression was reduced on CNS-infiltrating, but not peripheral, CD4+ T cells during EAE onset in Cav-1^-/-^ (Fig. 5C-D). Monocyte infiltration into the spinal cord but not the brain was reduced in Cav-1^-/-^ compared to WT mice at EAE onset (Fig. S2B). Nonetheless, there was no significant change in EAE disease severity at 10 DPI in Cav-1^-/-^ mice compared to WT (Fig. S2A). Cav-1 deficiency did not significantly alter total or CXCR3+ neutrophil or monocyte infiltration into the CNS or their peripheral abundance during EAE onset (Fig. S3A-D). We also analyzed a subset of chemokine receptors associated with Th17, regulatory T cell, and monocyte populations: CCR6, CXCR4, and CCR2 (Heng et al., 2022). We found there was no change in expression of these chemokine receptors on peripheral or CNS infiltrating leukocytes associated with Cav-1 deficiency during EAE onset (Fig. S3E-F).

**Figure 5:**
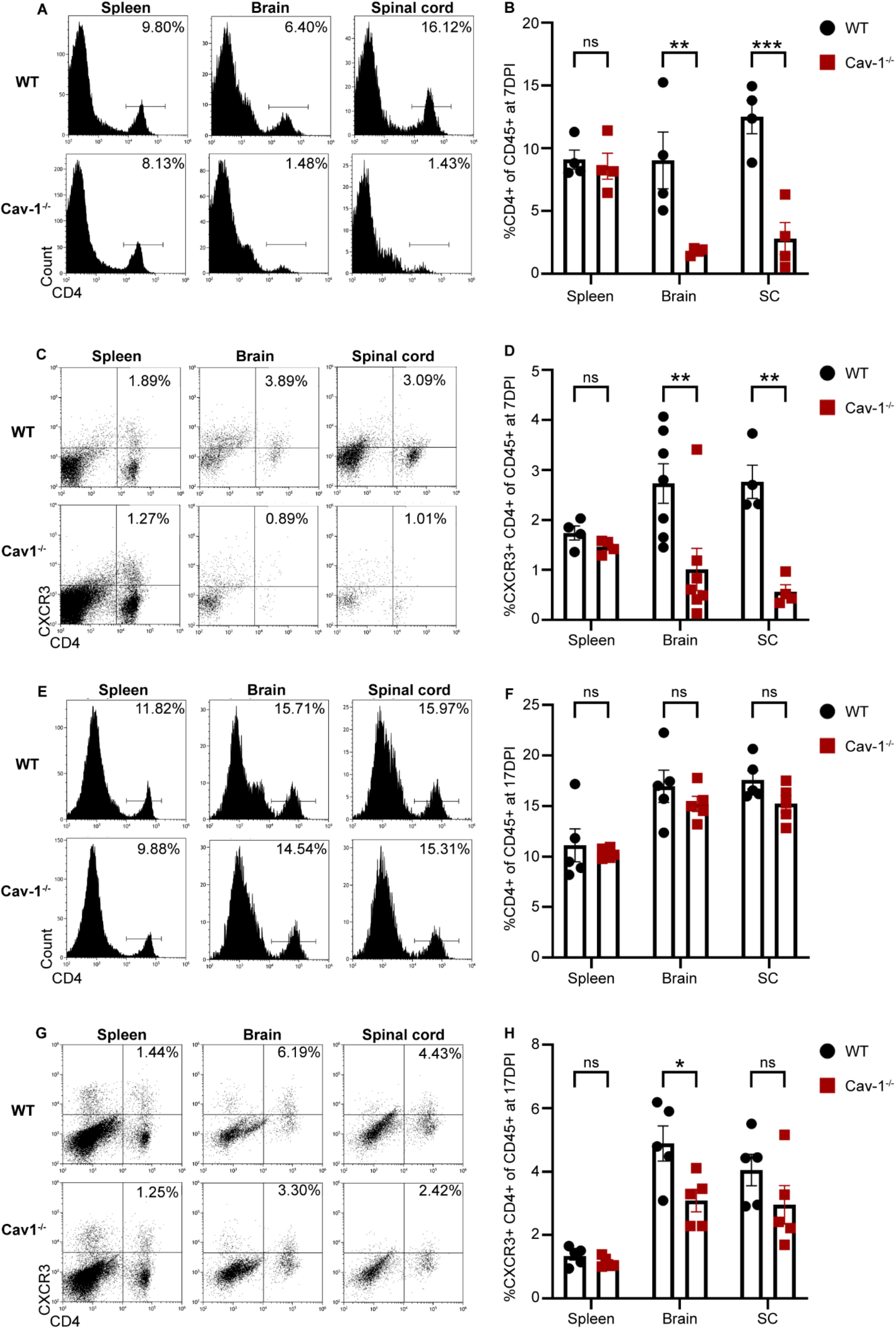
Cav-1 deficiency correlates with reduced CXCR3+ T cell CNS infiltration during EAE onset. **(A)** Representative histograms of flow cytometric analysis of CD4+ cells in spleen, brain, and spinal cord of WT and Cav-1^-/-^ mice during EAE onset (10 DPI) gated on viable, CD45+ cells. **(B)** Quantification of CD4+ cells in spleen, brain, and spinal cord from WT and Cav-1^-/-^ mice during EAE onset. **(C)** Representative dot plots of flow cytometric analysis of CXCR3+ CD4+ population of viable, CD45+ cells from spleen, brain, and spinal cord of WT and Cav-1^-/-^ mice during EAE onset. **(D)** Quantification of CXCR3+ CD4+ population of viable, CD45+ cells in flow cytometry analysis of spleen, brain, and spinal cord from WT and Cav-1^-/-^ mice during EAE onset. **(E)** Representative histograms of flow cytometry analysis of CD4+ population in spleen, brain, and spinal cord of WT and Cav-1^-/-^ mice during chronic EAE (17 DPI). **(F)** Quantification of CD4+ population of viable, CD45+ cells in flow cytometry analysis of spleen, brain, and spinal cord from WT and Cav-1^-/-^ mice during chronic EAE. **(G)** Representative plots of flow cytometry analysis of CXCR3+ CD4+ population of viable, CD45+ cells from spleen, brain, and spinal cord of WT and Cav-1^-/-^ mice during chronic EAE. **(H)** Quantification of CXCR3+ CD4+ population of viable, CD45+ cells in flow cytometry analysis of spleen, brain, and spinal cord from WT and Cav-1^-/-^ mice during chronic EAE. Two-way multiple comparison ANOVA with Sidak’s post-hoc comparisons between WT and Cav-1^-/-^ mice. * p<0.05 ** p>0.01 ***p<0.001

Disease severity continued to worsen until 17 DPI in both WT and Cav-1^-/-^ mice with EAE (Fig. 4A). At 17 DPI, we observed a significant reduction in disease severity in the Cav-1^-/-^ cohort (Fig. S2B). At this later time, there was no decrease in overall CD4+ T cell number in the CNS in Cav-1^-/-^ mice (Fig. 5E-F). The reduction of CXCR3+ T cells in the brain persisted in the Cav-1^-/-^ brain, but not the spinal cord, throughout the course of EAE (Fig. 5G-H). These data suggest that T cells infiltrating the CNS during the onset of disease utilize Cav-1 dependent transcellular migration to pass the BBB, whereas T cells can cross the BBB with or without endothelial Cav-1 expression in later stages of disease.

## Discussion

We used *in vitro* and *in vivo* approaches to elucidate the mechanisms whereby endothelial Cav-1 could facilitate T cell extravasation during the early stages of neuroinflammation. We found that CXCL10 and Cav-1 interacted to enhance transcellular but not paracellular migration of T cells across primary BBB ECs. First, endothelial cells presented CXCL10 in chemokine depots that were regulated in size and stability by Cav-1 expression. Second, CXCL10 primed CXCR3+ CD4+ T cell for transcellular diapedesis through brain endothelial cells, which was also dependent on endothelial Cav-1 expression. These data were supported and extended by *in vivo* studies that established endothelial cell Cav-1 expression enhances CNS infiltration of CXCR3+ T cells, using adoptive transfer studies and endothelial cell conditional knockout of Cav-1. The work presented herein highlights two distinct phases of T cell BBB extravasation *in vivo*: an initial phase in which T cell entry into the brain is enhanced by endothelial Cav-1 regulation of processes including chemokine presentation and transcellular permeability, and a later stage coinciding with fulminant disease where Cav-1 regulated processes are dispensable. Overall, our data support a mechanistic pathway in which endothelial CXCL10 activates T cell CXCR3, promotes inside-out activation of T cell LFA-1, enhances LFA-1 binding to endothelial ICAM clusters organized in the membrane by Cav-1, and causes transcellular transmigration of autoreactive CD4+ T cells across the BBB at the onset of disease (Fig 6). This data contributes to the emerging appreciation of dynamic phases of BBB permeability in time and space, with implications for immune surveillance, neuroinflammation, and disease recovery (Liebner et al., 2018; Marchetti & Engelhardt, 2020).

**Figure 6:**
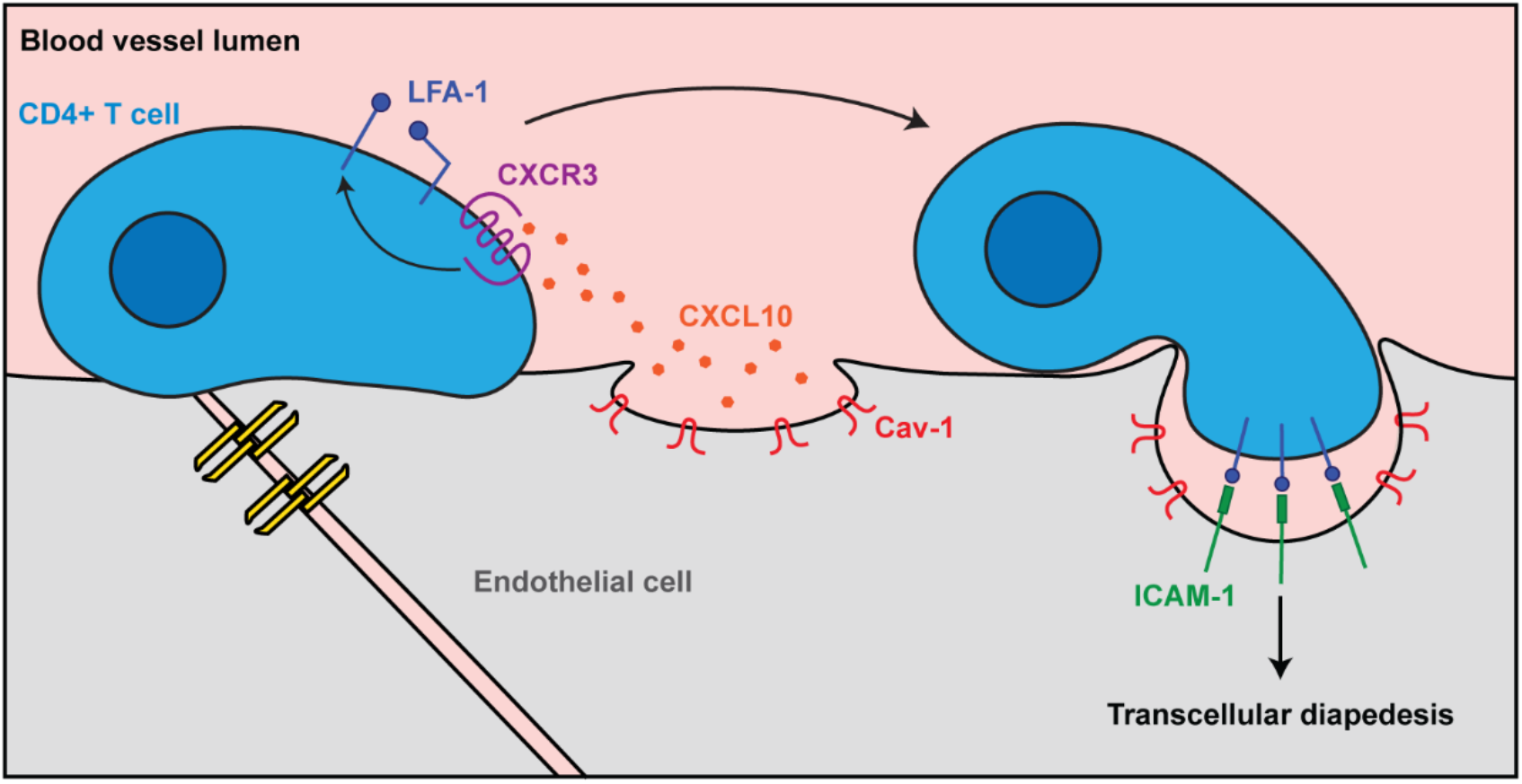
CXCR3 and Cav-1 induce transcellular diapedesis of T cells at the BBB. CXCR3+ CD4+ T cells are activated by brain endothelial cell CXCL10 depots formed with Cav-1. CXCR3 signaling activates integrin LFA-1. LFA-1/ICAM-1 interaction facilitates Cav-1-dependent transcellular diapedesis of CXCR3+ CD4+ T cells across the BBB.

Our findings advance our understanding of how T cells are recruited for transcellular, instead of paracellular, transmigration. We found that transcellular but not paracellular migration involved CXCL10, Cav-1, and LFA-1 activity. Endothelial cells in an inflammatory milieu promote transcellular permeability in part by aggregating chemokines and adhesion molecules into focal pockets that support transmigration (Carman & Springer, 2008; Shulman et al., 2011). We found that in response to IFNγ, brain endothelial cells expand their intracellular chemokine stores. In the absence of Cav-1 expression, endothelial cells instead secreted these chemokines. The depletion of intracellular chemokine stores in the absence of Cav-1 is likely to reduce T cell diapedesis, because local chemokine release activates T cell chemokine receptors and integrins such as LFA-1. LFA-1 binding crosslinks and activates its cognate cell adhesion molecule ICAM-1 on endothelial cells (Ngwenyama et al., 2019; Prizant et al., 2021). In this process, ICAM-1, F-actin, and Cav-1 closely associate in docking structures in the EC membrane (Barreiro et al., 2002; Carman & Springer, 2004; Heemskerk et al., 2016; van Buul et al., 2007). LFA-1 binding of ICAM-1 activates Cav-1 phosphorylation by Src and promotes transcellular hyperpermeability (Liu et al., 2011). Endothelial ICAM-1 ligation and clustering also induced regulated release of vesicular stores of CXCL1, CXCL8, and CXCL10 and promoted transcellular diapedesis (Schoppmeyer et al., 2022). As much as 90% of endothelial CXCL10 is sequestered in intracellular stores, with release regulated by synaptic exocytosis machinery (Schoppmeyer et al., 2022). Thus, Cav-1 may promote transcellular T cell diapedesis by enabling the formation of inflammation-induced vesicular chemokine compartments that provide chemokines directly to T cell invadopodia-like protrusions.

We have not established how Cav-1 controls the expansion or secretion of chemokine pools. Interestingly, however, Cav-1 is well-characterized for its structural role in caveolar vesicles, and phosphorylation of Cav-1 Tyr14 promotes a space-filling conformational change that increases the volume and motility of Cav-1 intracellular vesicles loaded with macromolecular cargo (Zimnicka et al., 2016). Furthermore, Cav-1 colocalizes with ACKR decoy chemokine receptors known to promote vesicular delivery of chemokines (Cruz-Orengo et al., 2011; Girbl et al., 2018; Marchetti et al., 2022; Minten et al., 2014), suggesting that Cav-1 might contribute to the intracellular trafficking of chemokines by ACKR interactions (Cruz-Orengo et al., 2014; Luo et al., 1997; Middleton et al., 1997; Pruenster et al., 2009). Indeed, ACKR1 specifically enhances transcellular T cell diapedesis across brain EC (Marchetti et al., 2022; Minten et al., 2014), and depleting intracellular chemokine depots by genetic deletion of ACKR1/ACKR3 in brain EC decreased EAE severity — due to decreased T cell recruitment across the BBB (Cruz-Orengo et al., 2011; Minten et al., 2014).

Strategies to manipulate CXCL10 in neuroinflammation have had mixed outcomes (Heng et al., 2022; Karin, 2020). For example, astrocyte-targeted CXCL10 depletion lessened EAE severity by reducing perivascular accumulation of effector T cells and acute demyelination, whereas global deletion of CXCL10 did not protect against disease (Klein et al., 2004; Mills Ko et al., 2014). Several studies did show EAE reduction upon deletion or blockade of CXCL10 or CXCR3 (Kohler et al., 2008; Sporici & Issekutz, 2010). Our data suggest that endothelial Cav-1 promotes transcellular but not paracellular migration of CXCR3+ T cells across the BBB. Notably, however, we found that CXCR3^-/-^ T cells were still able to diapedese using the paracellular route. This is consistent with a previous report that CXCL10/CXCR3 deficiency does not prevent CNS entry of adoptively transferred MOG-specific Th1 cells (Lalor & Segal, 2013). Interpretations are complicated because CXCL10/CXCR3 signaling has functions in multiple cellular compartments, during different pathogenic events, and as part of a network of interrelated and partially compensatory chemokine signaling loops (Duckworth et al., 2021; Karin, 2020; Koper et al., 2018). Because CXCL10 is transcriptionally regulated by IFNγ, the mechanistic pathway we describe in the present paper might be most relevant for neuroinflammation in infectious diseases where interferon production is high (Blank et al., 2016; Coperchini et al., 2021; Olivarria et al., 2022; Sorensen et al., 2018). Inflammation also increases BEC Cav-1 expression and activity (Hu et al., 2008; Lutz et al., 2017; Wu et al., 2016), which can also increase CNS exposure to infectious agents (Salimi et al., 2020). Our studies provide a mechanistic basis for why previous efforts to harness CXCR3 chemokine receptor antagonism to control MS/EAE have had inconsistent and ultimately unsuccessful results (Heng et al., 2022; Muller et al., 2010).

Signals from the BEC to the T cell during diapedesis can have lasting effects on T cell behavior within the target tissue, after diapedesis is complete. For example, the heightened LFA1/ICAM1 interactions associated with transcellular diapedesis are expected to promote proliferation, enhancement of effector functions, and T cell memory development (Gerard et al., 2021). We found that the effect of endothelial cell Cav-1 in CNS infiltration of CXCR3+ T cells was most pronounced during disease onset. Therefore, CXCR3+ T cells encountering LFA-1 during transcellular transmigration in the early phases of lesion formation are expected to produce more IL-17, IFNγ, GMCSF, and other signals that activate endothelial cells and glia to further disrupt BBB integrity (Dusi et al., 2019; Kebir et al., 2007; Platt et al., 2020). Schoppmeyer and colleagues found T-cell binding stimulated endothelial cells to simultaneously extrude CXCL1, CXCL8, and CXCL10 (Schoppmeyer et al., 2022). Combinatorial chemokine exposure potently increases blood-cerebrospinal fluid barrier migration (Schlager et al., 2016). Therefore, it appears likely that transcytosis heightens T cell activation by triggering multiple chemokine receptor signaling cascades. Furthermore, endothelial ICAM-1/2 and other costimulatory receptors encountered during diapedesis can activate or inhibit differentiation, cytokine expression, and proliferation of T cells (Manes et al., 2022). It remains to be determined how BEC chemokine, integrin, and costimulatory signals activated during crawling and transcellular or paracellular diapedesis influence immune surveillance and neuroinflammation (Heng et al., 2022; Manes et al., 2022; Mapunda et al., 2022). Together, these studies suggest that transcellular diapedesis could be important for the recruitment of highly activated pioneer T cells that participate in the early phases of neuroinflammation. This work clarifies the mechanism by which Cav-1 facilitates transcellular permeability and suggests new avenues for altering the CNS infiltration of autoreactive or protective immune cells in neuroinflammatory diseases.

## Acknowledgements

We thank Ms. Maricela Castellon for expert assistance with mouse husbandry. This work was supported by funds from the University of Illinois at Chicago Department of Anatomy and Cell Biology and KL2TR002002 (SEL), T32 HL144459 (TNT), R01 AG061114 (LMT).

## Supplemental Figures

**Figure S1:**
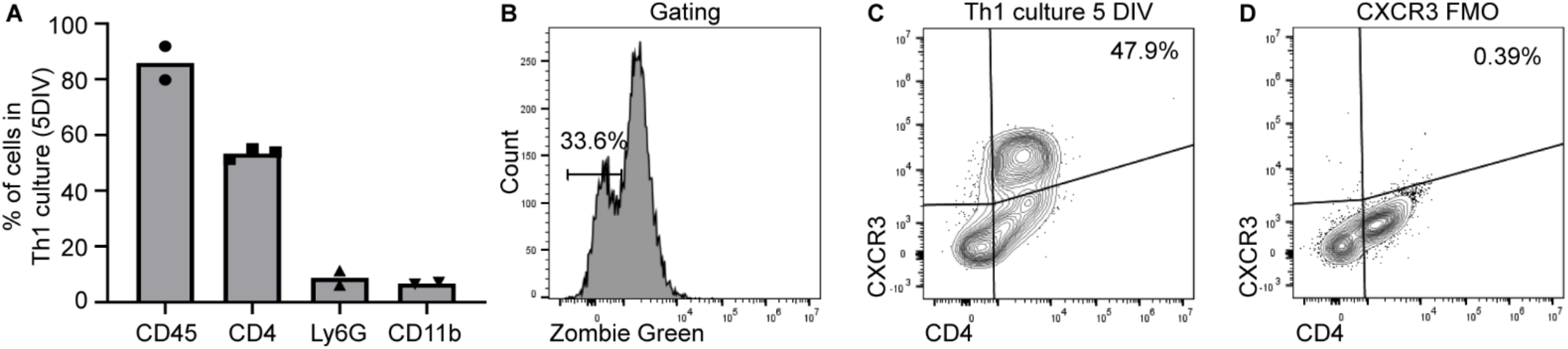
CXCR3+ CD4+ T cell enrichment in culture. **(A)** Immune cell composition of total culture composition after 5 days *in vitro* (DIV) by immunofluorescence. **(B)** Gating strategy to exclude nonviable cells (Zombie Green+). **(C)** Viable CXCR3+ CD4+ T cells were assessed from Th1 cultures at 5 DIV. **(D)** CXCR3 antibody was omitted for the fluorescence minus one (FMO) control condition.

**Figure S2:**
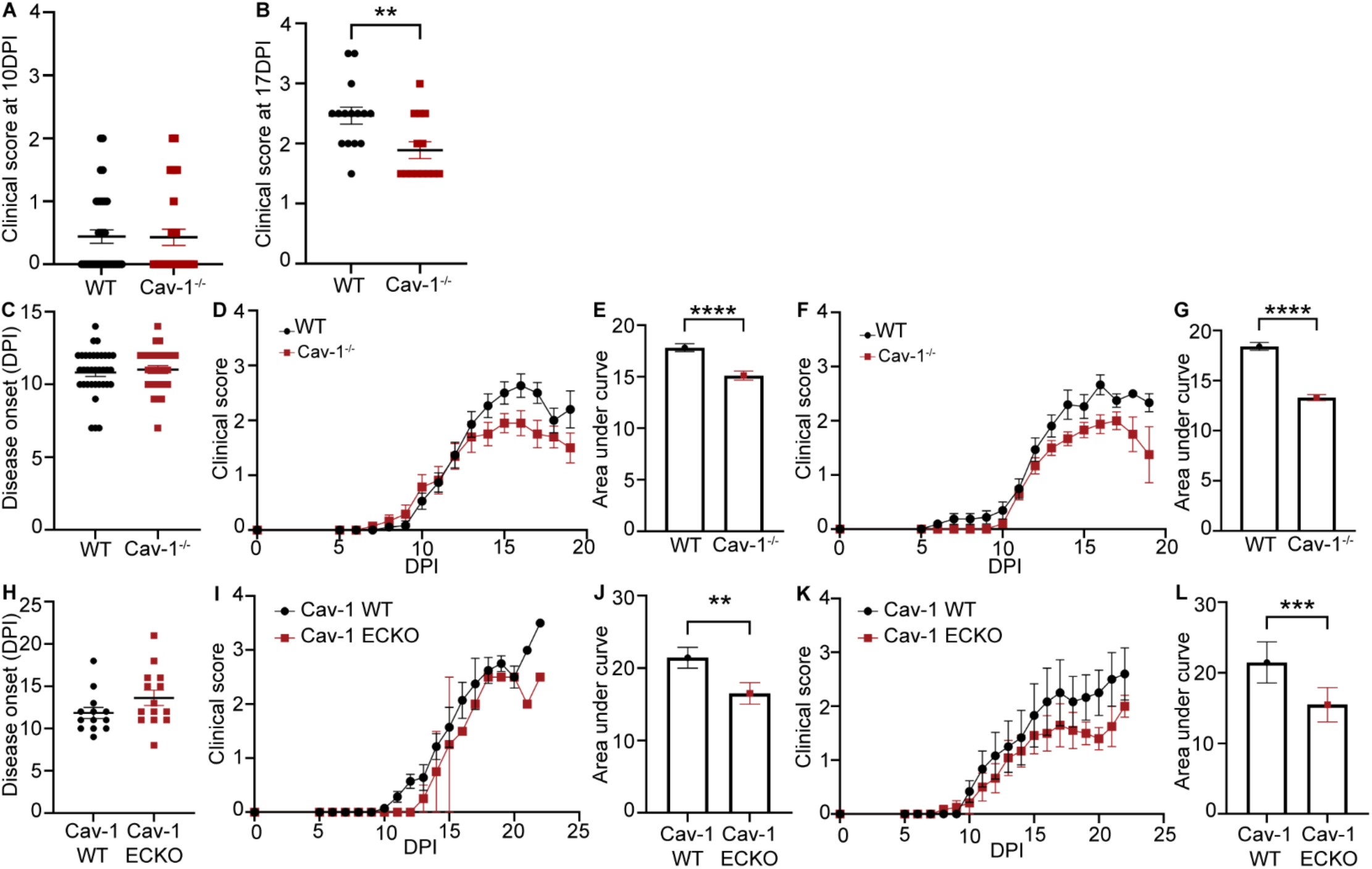
EAE disease onset and sex separated clinical curves. **(A)** Clinical scores of WT and Cav-1^-/-^ EAE mice at 10 DPI. **(B)** Clinical scores of WT and Cav-1^-/-^ EAE mice at 17 DPI. **(C)** Date of onset of clinical signs for WT and Cav-1^-/-^ EAE. **(D)** Clinical curve of female WT (n=18) and Cav-1^-/-^ EAE (n=14). **(E)** Area under the curve analysis of female WT and Cav-1^-/-^ clinical curves 0-19 DPI. **F**. Clinical curve of male WT and Cav-1^-/-^ EAE. **(G)**. Area under the curve analysis of male WT (n=16) and Cav-1^-/-^ (n=15) clinical curves 0-19 DPI. **(H)** Disease onset of Cav-1 WT and Cav-1 ECKO EAE. **(I)** Clinical curve of female WT (n=7) and Cav-1^-/-^ (n=2) EAE. **(J)** Area under the curve analysis of female WT and Cav-1^-/-^ clinical curves 0-22 DPI. **(K)** Clinical curve of male WT (n=6) and Cav-1^-/-^ (n-12) EAE. **(L)** Area under the curve analysis of male WT and Cav-1^-/-^ clinical curves 0-22 DPI. Unpaired t-test ** p<0.01 *** p<0.001 **** p<0.0001

**Figure S3:**
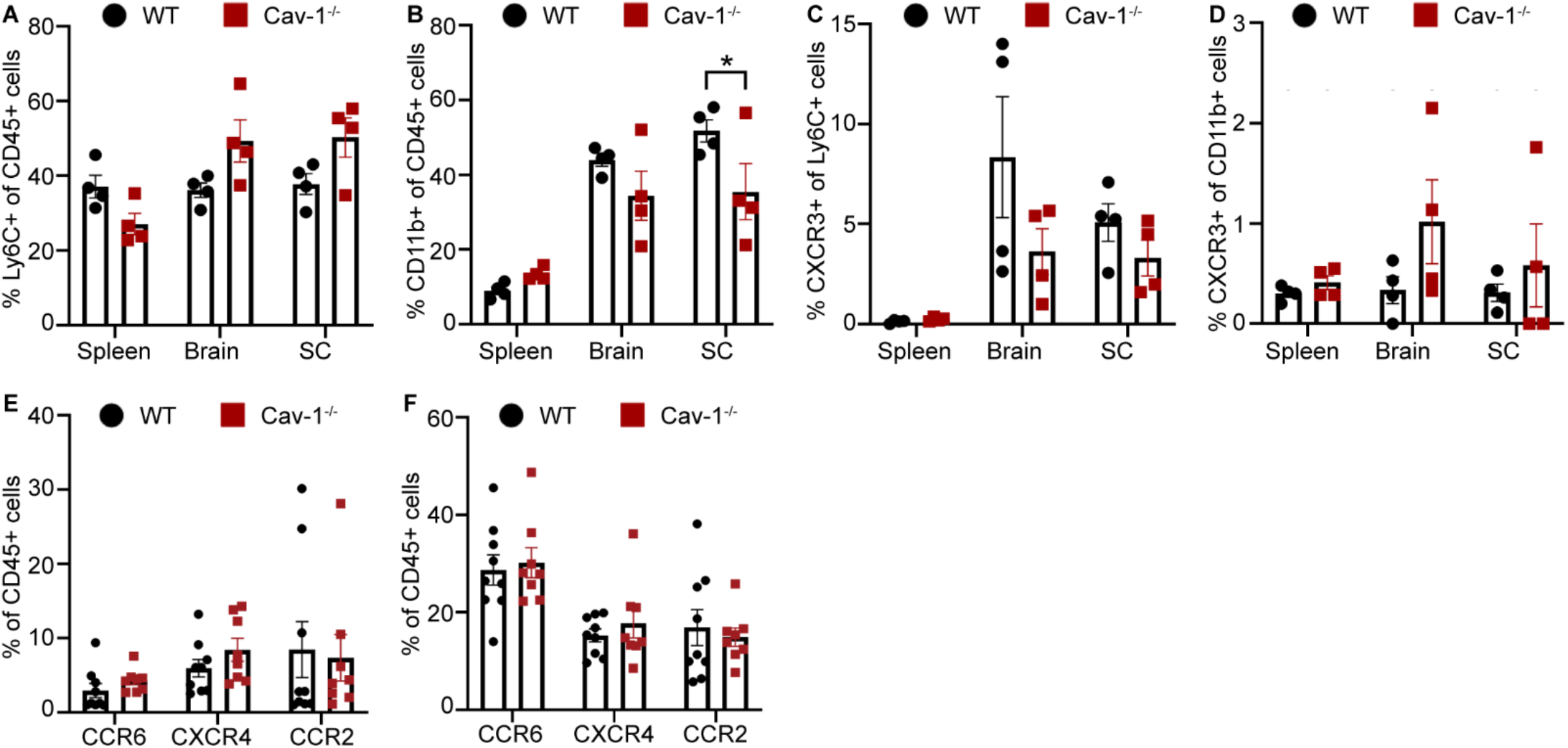
Flow cytometric analysis of innate immune cells and chemokine receptor expression in EAE. **(A)** Quantification of Ly6C+ population of viable, CD45+ cells in flow cytometric analysis of spleen, brain, and spinal cord from WT and Cav-1^-/-^ mice during EAE onset at 10 DPI. **(B)** Quantification of CD11b+ population of viable, CD45+ cells in flow cytometry analysis of spleen, brain, and spinal cord from WT and Cav-1^-/-^ mice during EAE onset. **C**. Quantification of CXCR3+ Ly6C+ population of viable, CD45+ cells in flow cytometry analysis of spleen, brain, and spinal cord from WT and Cav-1^-/-^ mice during EAE onset. **(D)** Quantification of CXCR3+ CD11b+ population of viable, CD45+ cells in flow cytometry analysis of spleen, brain, and spinal cord from WT and Cav-1^-/-^ mice during EAE onset. **(E)** Chemokine receptor analysis of CD45+ cells from spleen of WT and Cav-1^-/-^ mice during EAE onset. **(F)** CCR6, CXCR4, and CCR2 chemokine receptor positive CNS-infiltrating leukocytes from WT and Cav-1^-/-^ mice during EAE onset. Two-way ANOVA with Sidak’s post-hoc comparisons between WT and Cav-1^-/-^ groups. * p<0.05

